# Two new threatened *Inversodicraea* (Podostemaceae) species from Sierra Leone: *I. joulei* and *I. lebbiei*

**DOI:** 10.64898/2026.05.18.725858

**Authors:** Fanny Massally, Aiah Lebbie, Xander M. van der Burgt, Jack Plummer, Martin Cheek

## Abstract

Two threatened new species of Podostemaceae belonging to the genus *Inversodicraea, I. joulei* and *I. lebbiei*, both from the Republic of Sierra Leone, are described and illustrated. A first record in Sierra Leone of the genus *Lestestuella* is also reported. *Inversodicraea* is the most species-rich genus of Podostemaceae in Africa and now comprises 38 species. *Inversodicraea joulei* is easily recognised because it has a persistent spine distally on the median rib of each fruit valve, and scattered, membranous scale-leaves with broadly rounded apices, while *Inversodicraea lebbiei* is distinct in having narrowly triangular robust scale-leaves which are inrolled, spreading distally, and completely covering the stem, arranged in five ranks. *Inversodicraea joulei* is known from a single location with three sites while *I. lebbiei* is known from two locations each with one site. Using the latest IUCN Red List guidance, *Inversodicraea joulei* is assessed as Critically Endangered and *I. lebbiei* is assessed as Endangered, due to threats from dam construction projects, agricultural practices and mining activities, resulting in high levels of siltation on rocks in the fast-flowing rivers where these species grow.

## Introduction

Podostemaceae Rich ex Kunth, also known as the “Orchids of the falls,” represents one of the most ecologically specialized and morphologically unique families (Philbrick and Novelo 1995, Rutishauser, 1995). These rheophytic herbs exhibit extraordinary evolutionary adaptations that enable them to thrive in the challenging conditions of fast-flowing, clear-water rapids, and waterfalls, allowing them to colonize seasonally submerged rocks where few other vascular plants can endure (Rutishauser et al 1999; Koi *et al.et al*. 2012; Kato, 2016; Cheek *et al.et al*. 2020). There are 52 accepted genera in the Podostemaceae family (POWO 2025). Species numbers are highest in tropical America, followed by Asia, with Africa having c. 80 species (Cheek *et al*. 2017a). Many African Podostemaceae species are narrow endemics, with some restricted to a single river system or waterfall, making them particularly susceptible to habitat degradation (Rutishauser, 2004).

*Inversodicraea* Engl. is Africa’s most species-diverse Podostemaceae genus (Cheek *et al*. 2020). *Inversodicraea* is placed in the subfamily Podostemoideae and has undergone significant taxonomic re-evaluation over recent decades. Initially, species now placed in *Inversodicraea* were classified under *Ledermanniella* subgenus *Phyllosma* C. Cusset (1973, 1984, 1987). However, Thiv *et al*. (2009) resurrected the genus to include all African Podostemoideae taxa with scale-leaves, distinguishing them as a discrete subclade based on morphological and molecular evidence. A subsequent synoptic revision by Cheek *et al*. (2017a) transferred additional species to and recognized 30 species of *Inversodicraea*, distinguished from *Ledermanniella* by dorsiventrally flattened, spirally arranged scale-leaves usually densely covering the stems. Since then, several more new species of *Inversodicraea* have been described, all of which are highly range-restricted and Critically Endangered (Cheek *et al*. 2019; 2020). Sierra Leone is divided into four broad topographic regions, namely, the coastal lowland, the interior plains, the interior plateau and scattered mountains and hills. There are nine major rivers systems that are a host to a mosaic of Podostemaceae habitats and other aquatic biomes (Norman et al 2018, Fayiah *et al*. 2020). Recent botanical explorations have highlighted the country’s remarkable potential for hosting endemic rheophytic flora. Two decisive moments in this context were the discovery of *Lebbiea grandiflora* Cheek, a new genus and species of Podostemaceae in the rapids of the Sewa River (Cheek & Lebbie 2018), as well as *Ledermanniella yiben* Cheek from the Seli River (Cheek *et al*. 2017a). These findings indicate the uniqueness of Sierra Leone’s aquatic habitats and the potential for further discovery of other unknown taxa. However, the fragile habitats in which these plants occur face imminent threats. Some of Sierra Leone’s waterfalls and rapids, the primary habitats for Podostemaceae, are scheduled for hydroelectric development. Increased anthropogenic activities in areas where these plants grow, such as artisanal gold mining, could lead to irreversible species loss before being documented.

Following the discovery of *Ledermanniella yiben* (Podostemaceae) in the Seli River and recognizing the imminent threats to its survival, further surveys were recommended of the rapids and waterfalls in Sierra Leone. The objective was to discover, if possible, additional locations for this species. Accordingly, in 2017 and 2018, Aiah Lebbie and his team collected 70 Podostemaceae specimens from various river systems in Sierra Leone, including the Little Scarcies, Kaba River, Sewa River, Bagbe River, Rokel River, and River Waanje (Lebbie, 2017, Map 1). One of these specimens was determined to represent a new genus and described as *Lebbiea grandiflora* (Cheek & Lebbie 2018). Further study of these 70 Podostemaceae specimens resulted in the discovery of two additional new species. This publication identifies and describes two new species of *Inversodicraea*, presenting a taxonomic key to the species of *Inversodicraea* found in Sierra Leone. It was completed as part of an MSc project by the first author (Massally 2025).

## Materials and Methods

Field observations and herbarium collections of *Inversodicraea* were made in Sierra Leone by two of the authors, Aiah Lebbie and Fanny Massally. All studied specimens are kept at the National Herbarium of Sierra Leone (SL) with duplicates at the Kew Herbarium (K). Specimens from the National History Museum (BM) were also checked. Herbarium collections were examined with a Leica S6E microscope with a calibrated eyepiece graticule measuring objects of 0.025 mm or more at maximum magnification. Additionally, some macroscopic parts were measured using a ruler. The drawings (Figs. 1 and 2) were made using a Leica 308700 camera lucida attachment. The specimens from the survey of Lebbie (2017) were mainly collected in April and May 2017 at the end of the dry season, when most Podostemaceae, including those described in this paper, have already fruited and died. Thus, the material was not in the ideal state for study. Nevertheless, in most cases, floral structures were preserved and described. All names of species and authors are based on the International Plant Names Index (IPNI continuously updated) website. The descriptions adhere to the format established by Cheek *et al*. (2017a). The conservation assessment follows current IUCN standards, with a grid cell size of 4 km^2^ (IUCN 2012; 2024). A distribution map for the two new species described (Map 2) was created using Simple Mappr (http://www.simplemappr.net/ ).

**Fig. 1.**
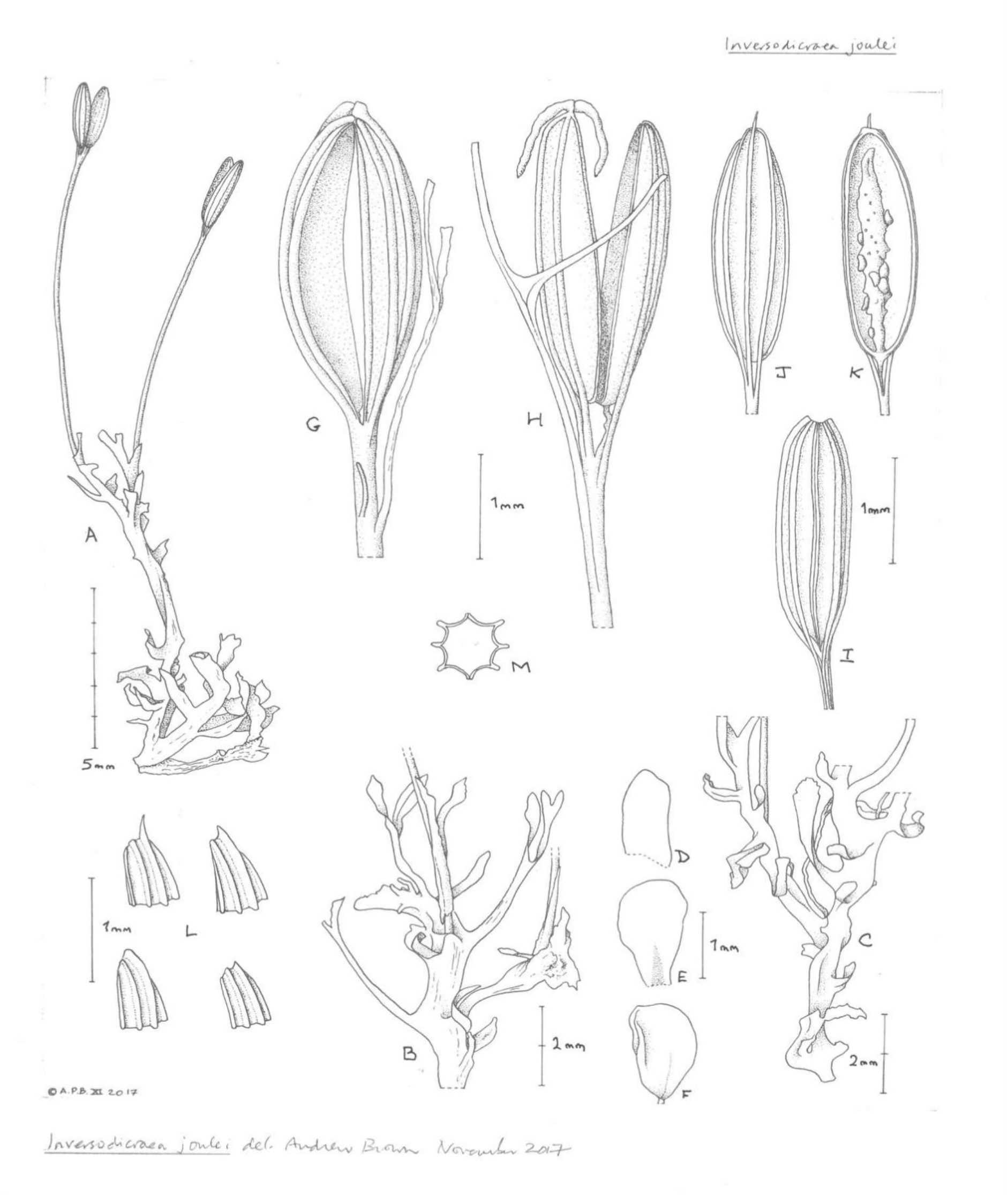
*Inversodicraea joulei*. Cheek & Massally. A. habit, branched stem with fruits; B. stems with dichotomous ribbon leaves; C. stems with sparse, entire scale-leaves; D-F. scale-leaves from B & C; G. fruit with two persistent staminal filaments (anthers missing) and filiform tepal (hydrated); H. dehisced fruit with two persistent stigmas (hydrated); I. undehisced fruit: note the well-developed commissural ribs (dry state); J. exterior fruit valve with long median “spine” (dry state); K. dehisced fruit with seeds attached to placenta (1 valve removed); L. heteromorphic median spines (dry state); M. transverse section of the fruit (A-F, J-M from *Lebbie 2690*; H from *Lebbie 2694*; G from *Lebbie 2696*; all K). Drawn by Andrew Brown.

**Fig. 2.**
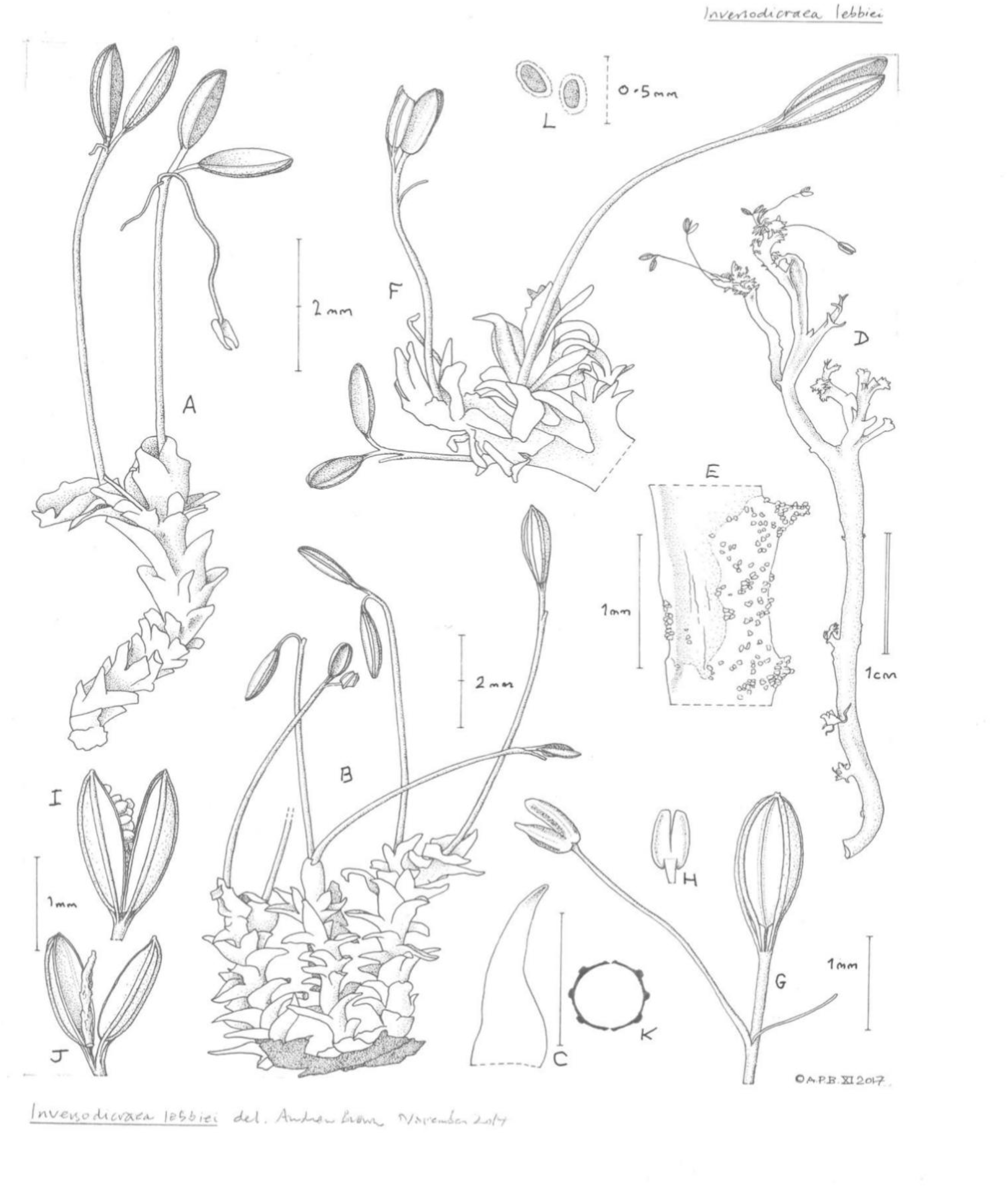
*Inversodicraea lebbiei*. Cheek & Massally. A. habit, stems with scale-leaves, dehisced fruits with tepal and stamen persisting (Scale-leaves somewhat degraded); B. habit, stems on remnant of root attached to rock; C. scale leaf from B; D. ribbon-like root with paired marginal shoots; E. crystals on root surface; F. from D, shoots with dehisced and undehisced fruits (note the longer scale-leaves towards the apex); G. fruit with persisting flower parts (note the barely developed commissural ribs of the fruit); H. anther, dorsal view; I. dehisced fruit with seeds; J. dehisced fruit showing placenta; K. transverse section of fruit; L. hydrated seed with mucilage coat (A, D, E, G, & H from *Lebbie 2709* and B, C, F, & I-L from *Lebbie 2726*; all K). ― Drawn by Andrew Brown.

**Map 1:**
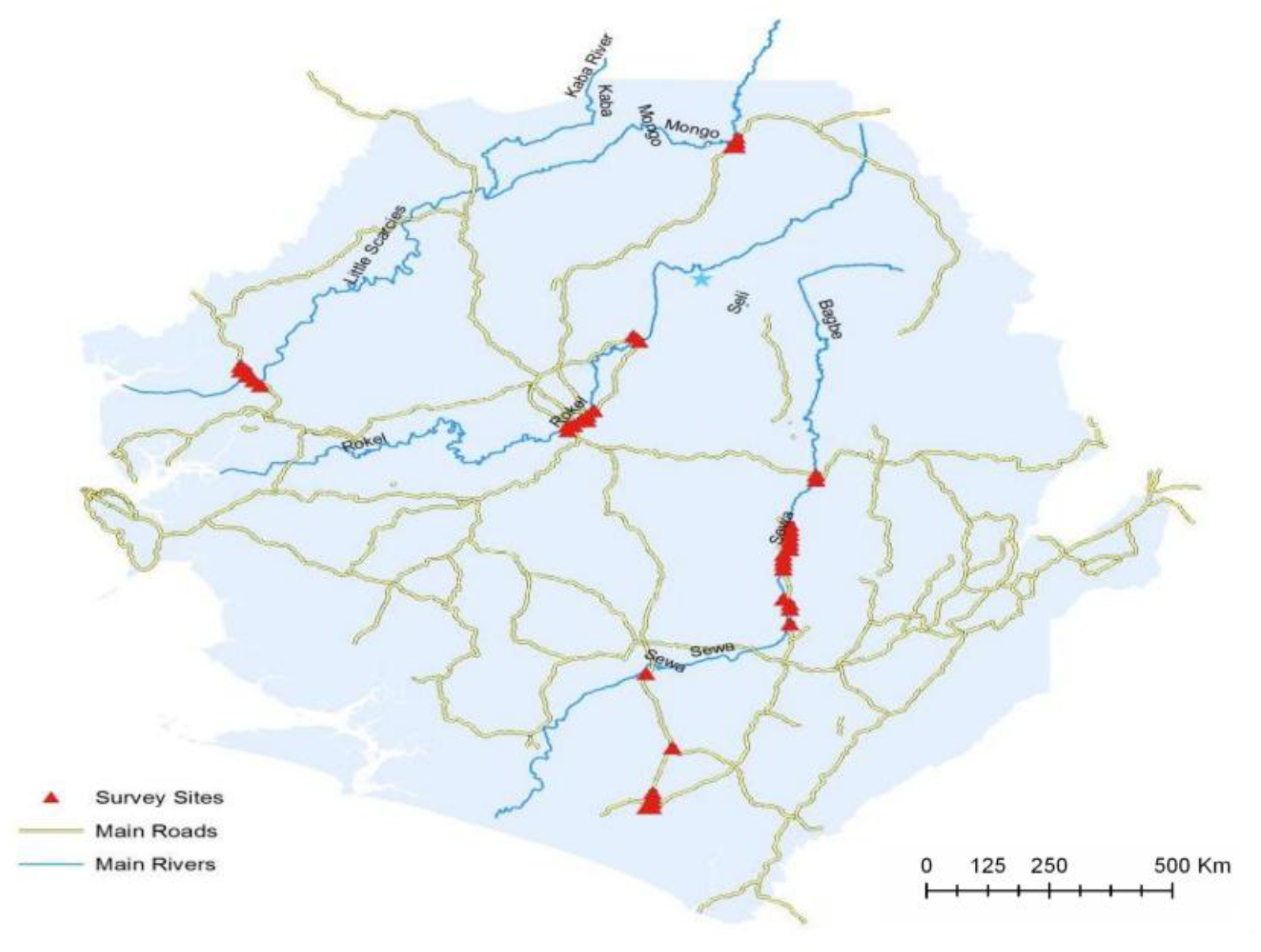
Map of river sites surveyed for Podostemaceae by the Lebbie team in 2017 (from Lebbie, 2017).

**Map 2:**
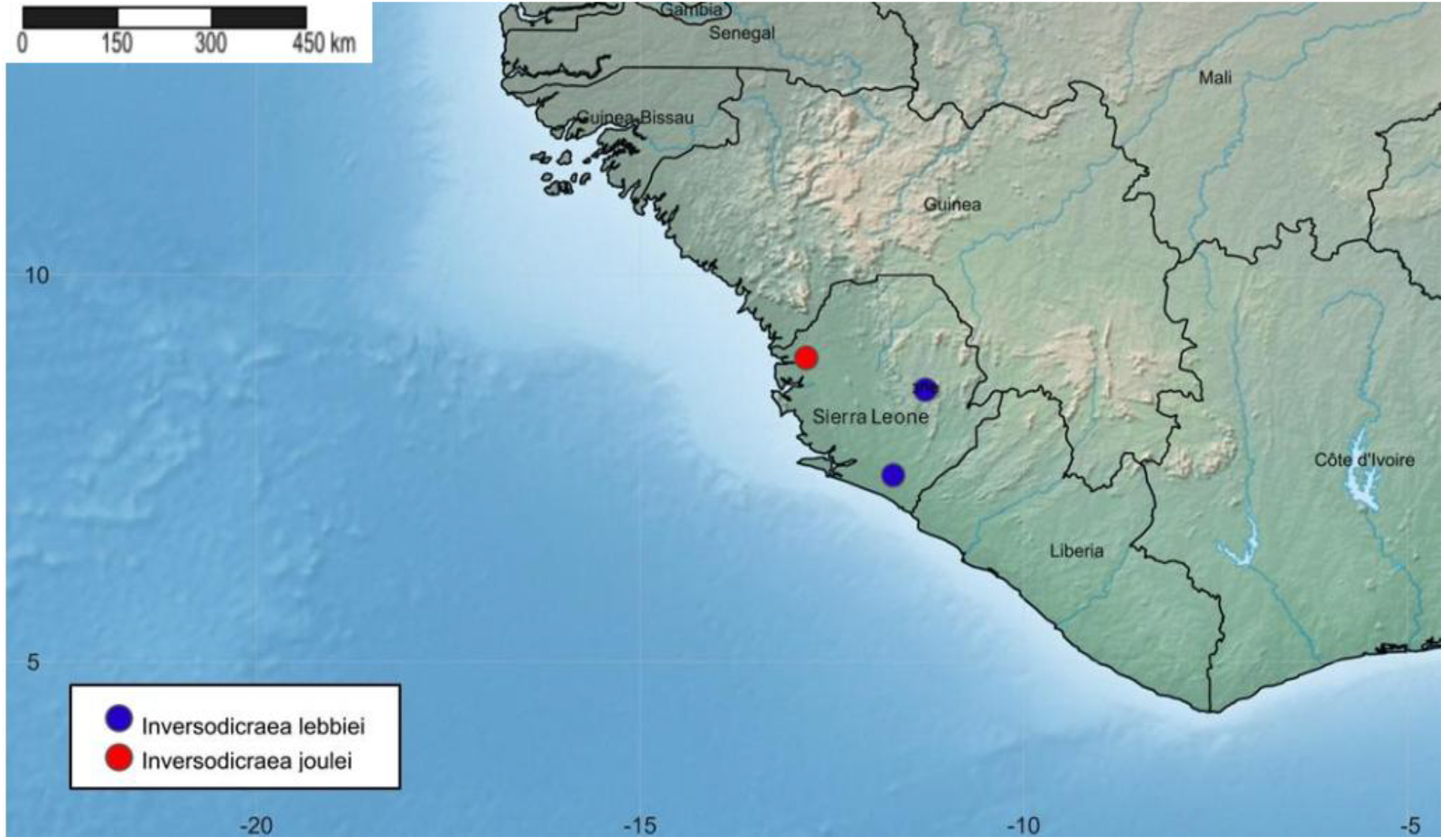
Global distribution map of *Inversodicraea joulei* and *I. lebbiei* in Sierra Leone.

## Results

### Key to species of *Inversodicraea* in Sierra Leone (modified from Cheek *et al*. 2017a)

1. Scale-leaves densely covering (≥ 50%) both the principal stems and the side branches (if present) ……………………………………………………………………………………………………………………..2
1. Scale-leaves present only on the side branches and/or sparsely covering (< 40%) the principal stems …………………………………………………………………………………6
2. Scale-leaves entire or longer than wide, with 0 – 2 lateral lobes ……………………………3
2. Scale-leaves distinctly 3( – 4)-lobed …………………………………………………………5
3. Scale-leaves linear-subulate ………………………………………………………***I. mortonii***
3. Scale-leaves ovate, lanceolate, ligulate, orbicular, or obovate; apex dentate, lobed, or rounded ………………………………………………………………………………………..4
4. Apex of leaflet lobes ±acute; 1 stamen; commissural ribs barely developed ***I. lebbiei sp*.*nov***.
4. Apex of leaflet lobes broadly rounded and rarely acute; 2 stamens; commissural ribs weakly developed …………………………………………………………………..***I*.*cf. liberia***
5. Scale-leaves concave, three apical lobes curved inward, with a fourth reflexed lobe on the abaxial surface; fruit with 6 longitudinal ribs …………………………………………***I. harrisii***
5. Scale-leaves flat with three ± equal apical triangular lobes but lacking lobe on abaxial surface; fruit with 8 longitudinal ribs …………………………………………………………………………………………………………………………………………………***I. ledermannii***
6. Scale-leaves with apex ± rounded; fruit with commissural ribs well-developed; median fruit ribs with distal ‘spines’ ………………………………………………………..***I. joulei sp*.*nov***.
6. Scale-leaves with apex acute or obtuse; fruit with commissural ribs absent or weakly developed; fruit ribs without spines ……………………………………………………………7
7. Scale-leaves c. 2 mm long, present on principle stems; androecium: gynoecium length c. 2:1 ……………………………………………………………………………………..***I. adamesii***
7. Scale-leaves < 0.5 mm long, present ± entirely on short spur-shoots subtending the spathellae; androecium: gynoecium length c.1:1 ………………………………………. ***I. feika***

**Notes**: *Lebbie* 2690A from Sierra Leone has scale-leaves in 3 ranks, covering 40 – 50% of the stem, and so is similar to *I. liberia* of Liberia. However, as the fruits and flowers are fragmentary, it is not described in this paper. It is provisionally named here as *I. cf. liberia*, **Inversodicraea joulei** *Cheek & Massally* **sp. nov**.

Type: Sierra Leone, Northern region, Port Loko District, near Bantoro and Mange villages, Little Scarcies River, 8° 55’ 29.7” N; 12° 50’ 58.0” W, alt. 15 m, fl., fr., 19 April 2017, *Lebbie* 2696 (holotype K [K001971291]; isotypes BR, MO, SL).

A rheophytic, probably annual herb, 13 – 85 mm tall (in fruit), vegetative parts drying black. ***Stems*** erect, drying flat, 0.2 – 1.3 mm diameter, 10 – 30% covered by scale-leaves, first branch 3 – 47 mm from the base, 2 – 3 subequal branches, each 2 – 35 mm long (Fig 1A). ***Roots*** not observed united to stems, fragmentary, ribbon-like, 0.75 – 1 mm wide, stems marginal at internodes of c. 1.5 mm apart. ***Leaves*** persistent, not falling at anthesis, bifurcate, with one possible occurrence of a double-bifurcate leaf (Fig 1B); leaf narrowly ribbon-like, arising laterally on stem, 3 – 5 × 0.2 – 0.3 mm, branching at 2 – 3 mm, base sessile, not sheathing, tapering gradually to obtuse apices. ***Scale-leaves*** simple, sessile, heteromorphic: obovate, elliptic, or oblong (Fig 1D – F), 0.5 – 3 × 0.2 – 0.5 mm, rounded, truncate to obtuse at apex. ***Spathella*** in flower bud not observed; in fruit single, cylindrical, 5 – 8 mm long, apex 0.2 – 0.8 mm wide, obtuse, base attenuate. ***Pedicel*** cream, 15 – 38 mm long in fruit. ***Tepals*** 2, caducous, inserted at base of gynophore (Fig 1G), filiform, 0.5 – 0.8 × 0.05 mm. Androphore and gynophore sub-opposite. ***Androecium*** with 2 stamens (Fig 1G & H); androphore when dried cream-coloured to black, ± 2 × 0.1 mm. Filaments 2, free, same colour as the androphore, c. 2 mm long. Anther and pollen not seen. ***Gynoecium*** with gynophore 0.6 – 1.4 × 0.2 – 0.3 mm (in fruit). ***Fruit*** a capsule (Fig 1I); cylindric-ellipsoid, slightly laterally compressed in transverse section (Fig 1M), 1.8 – 4 × 0.6 – 1 × 0.5 – 0.9 mm; ***stigmas*** 2, black when dried, fused at base, pendulous, botuliform, 0.6 – 1 × 0.1 mm, apex ± obtuse; Placentation free-central, ***placenta*** 1.5 – 3 mm long, 0.3 – 0.4 mm wide in the middle, tapering to the apex and base; capsule dehiscing into 2 equal valves. Valves persistent, inner surface glossy mid brown, externally dull brown flecked white; longitudinal ribs 6 (3 per valve), equal in length, highly pronounced, 0.05 – 0.1 mm wide, ribs decurrent into gynophore, in the dehisced fruit the 6 ribs exposed for c. 5 mm, connecting the valves and the pedicel (Fig. 1 J &K); commissural ribs 2 per valve, narrower than the longitudinal ribs, well developed but not as developed in depth and length, not extending to the apex of the fruit and the gynophore at the base (Fig 1J), 0.03 – 0.05 mm wide; median rib of each valve with spine at apex, median spine 0.1 – 0.2 mm long, obtuse to acute. ***Seeds*** ellipsoid, orange-brown, dark brown at both ends, 0.18 – 0.25 × 0.08 – 0.12 × 0.05 – 0.1 mm.

#### RECOGNITION

Morphologically similar to *Inversodicraea liberia* Cheek in having membranous scale-leaves, which are undulate when dry. However, it differs from *I. liberia* in having persistent distal spines on the median rib of the fruit valves (see Fig. 1J-L and Table 2). Additionally, the scale-leaves of *I. liberia* sometimes have a lateral triangular lobe (not entire as in *I. joulei*). Further, *I. liberia* capsules are also much smaller (1 – 1.1 × 0.5 – 0.8 mm) with weakly developed commissural ribs compared to *I. joulei’s* larger capsules (1.8 – 4 × 0.6 – 1 mm) and well-developed commissural ribs. Fig. 1, Map 2.

**Table 2:** Comparison of features used to distinguish *Inversodicraea joulei, I. lebbiei*, and *I. liberia* (data on the last species from Cheek *et al*. (2017a)

| Character | *I. joulei* | *I. lebbiei* | *I. liberia* |
| --- | --- | --- | --- |
| Scale-leaves apex shape | Broadly rounded | ± acute | Broadly rounded |
| Commissural ribs | Well developed | Barely developed | Only weakly developed |
| Fruit shape and width | Cylindric-ellipsoid, 0.6 – 1 mm | Ellipsoid, 0.5 – 0.8 mm | Ellipsoid, 0.6 – 0.7 mm |
| Scale leaf arrangement & shape | Irregularly arranged and obovate/elliptic/oblong | Arranged in c. 5 irregular ranks and narrowly triangular | Arranged in c. 3 ranks and obovate |
| Number of stamens per flower | 2 | 1 | 2 |
| Leaf branching | Twice dichotomous | Not recorded | Once dichotomous |
| Presence of spine on median rib of fruit | Yes | No | No |

#### SPECIMENS EXAMINED. SIERRA LEONE

Northern region, Port Loko District, Mange Bureh, Little Scarcies River, 8° 55’ 26.8” N; 12° 50’ 06.0” W, alt. 22 m, fl., fr., 17 April 2017, *Lebbie* 2690 (K [K001971289], P, SL, Z). Northern region, Port Loko District, near Bantoro and Mange villages, Little Scarcies River, 8° 56’ 12.7” N; 12° 50’ 44.3” W, alt. 24 m, fl., fr., 19 April 2017, *Lebbie* 2694 (B, K [K001971290], SL, WAG). Same locality, Little Scarcies River, 8° 55’ 29.7” N; 12° 50’ 58.0” W, alt. 15 m, fl., fr., 19 April 2017, *Lebbie* 2696 (holotype K [K001971291]; isotypes BR, MO, SL).

#### HABITAT

*Inversodicraea joulei* was collected on rocks in the Little Scarcies River, 15 – 24 m alt., with *Inversodicraea cf. liberia* and *Tristicha* species. These two species were found mixed with *Inversodicraea joulei* in the collection *Lebbie, A*. 2690 (K [K001971289]).

#### DISTRIBUTION

*Inversodicraea joulei* occurs in Sierra Leone, Northern Region, Port Loko District, Great Scarcies River, and is only known from three collections on this river made within 2 km distance from each other.

#### CONSERVATION STATUS

The collection sites for *Inversodicraea joulei* are under anthropogenic threat. Artisanal mining for gold, agricultural activities by communities upstream along the riverbank, and domestic activities, such as clothes washing, threaten the long-term survival of this species. The topography of the area is flat to slightly sloping; it does not seem to be a good site for the construction of a dam so likely this is not a threat. *Inversodicraea joulei* is known from three collections made in the same river, at less than 2 km apart from each other. The area of occupancy is 12 km^2^. The extent of occurrence, derived from herbarium specimen data and calculated using GeoCAT (Bachman *et al*. 2011), was measured as 1.075 km^2^. Because the actual extent of occurrence is less than the area of occupancy, the extent of occurrence should be changed to make it equal to the area of occupancy (IUCN 2024). The extent of occurrence is therefore 12 km^2^. The species does not occur further downstream because there are no more rapids as the river approaches the sea. The area of occupancy and the extent of occurrence are possibly larger than 12 km^2^. Based on the available information, the three collecting sites constitute one location, as they form a connected riverine system and the threats of gold mining, agricultural runoff and clothes washing could impact all three collection sites. There also exists a continuing decline in quality of habitat, and in the number of mature individuals. *Inversodicraea joulei* is here preliminarily assessed as Critically Endangered, CR B1ab(iii,v). This provisional assessment may encourage or guide further survey efforts prior to full assessment of all Podostemaceae in this river.

#### ETYMOLOGY

*Inversodicraea joulei* is named for the Joule Africa project, which funded the expedition that led to the collection of this species.

#### NOTES

*Inversodicraea joulei* is unusual in that the leaves, which in other species of the genus fall when plants are exposed by falling water levels and flower (Cheek *et al*. 2017a), are persistent, even at the fruiting stage. The specimens of the species were all collected at the beginning of the rainy season when the water level was rising. The material as a result was all long dead and dry when collected. It would be valuable to collect material when live and with complete flowers, and also roots, so that these might be characterised. We suggest that surveys should be targeted for the month of November to January.

The distinctive spine which arises from the distal median rib of the frut valves is developed after fruit dehiscence, when the valves become flat transversely, and only slightly curved along their length. The fruit spines, in combination with the black, sparsely scale-leaved, subshrubby stem, make this an easily recognised species.

The affinities of this species are possibly with those few other species in W Africa which have membranous leaves which dry with a wrinkled surface, these being *I. liberia* of Liberia and *I. pepehabai* Cheek of Guinea (Cheek & Haba 2016). Both those species however have the scale-leaves irregularly lobed and covering 50% more of the surface of the stem, they also lack the distinctive fruit spines of *I. joulei*.

### Inversodicraea lebbiei *Cheek & Massally* sp. nov

Type: Sierra Leone, Southern region, Kenema District, Sewa River, near Fomaya, Little Bekongor Falls, 8° 30’ 41.2” N; 11° 16’ 53.6” W, fl., fr., 243 m, 3 May 2017, *Lebbie* 2709 (holotype K [K001971287]; isotypes MO, P, SL, Z).

Rheophytic annual or possibly perennial herb, 6 – 75 mm tall (in fruit). ***Root*** horizontal, ribbon-like 0.6 – 1 (– 2) mm diameter, extending at least 4.5 cm (Fig 2D), with paired shoots (Fig 2F), from margin, internodes 4 – 9 mm long (Fig 2D). ***Leaves*** not seen. ***Stems*** either apparently annual, 6 – 12 mm tall, little branched; or possibly perennial, c. 5 cm long, with tough curved and branched stems 2 mm diam. at the base; erect, young stems covered by scale leaves, older stout, stems without scale-leaves. ***Scale-leaves*** covering the entire stem (except when older, thick stems are present) entire, robust, with numerous silica bodies, ascending, then reflexing, spreading, narrowly triangular, margins incurved, (Fig 2B & C), 0.8 – 1.4(– 2) × 0.2 – 0.5 mm, arranged in c.5 ranks along stem; scale-leaves elongated, clustered towards the apex (Fig 2F). ***Spathella*** in bud not seen, in fruit drying black, at dehiscence glabrous, 1 (– 2.5) × 0.5 – 0.6 (– 1) mm, stipe inconspicuous, apex opening by c. 3 longitudinal clefts half the length of the spathellum. ***Pedicel*** 4 – 7 mm long in fruit. ***Tepals*** 2, mostly fallen, filiform, inserted at the base of gynophore and androphore, 0.6 – 1 × 0.03 mm (Fig 2A & G).

Androphore and gynophore are inserted opposite each other (Fig 2G). ***Stamen*** monandrous (Fig 2A); filament transparent creamy brown, 2 – 3 × 0.05 mm. Anther (Fig 2G) bilobed, basifixed, creamy brown, 0.5 – 0.7 × 0.4 mm. ***Gynoecium*** with gynophore 0.5 – 1.2 × 0.1 mm (in fruit) and somewhat twisted. ***Fruit*** a capsule (Fig 2I & G), ellipsoid, 1.3 – 2 × 0.5 – 0.8 × 0.4 – 0.6 mm; hexagonal in transverse section, stigma probably bi-lobed, only a single lobe seen, staining black, coiled, 0.6 × 0.15 mm; placentation free, central, placenta 0.75 – 1 × 0.2 – 0.3 mm; capsule dehiscing into 2 sub-equal caducous valves outer surface brownish to yellow-brown, inner surface black-brown, apex obtuse, 0.05 – 0.1 mm long, base extending into gynophore; longitudinal ribs 6, equal in length, developed, decurrent, 0.05 mm wide; commissural ribs 2 per valve, narrower, barely developed, extending to apex and gynophore at the base (Fig 2G). ***Seeds*** with a mucilage coating when hydrated (Fig 2L), orange-yellow with darker tips at both ends, ellipsoid, 0.2 – 0.3 × 0.1 – 0.15 × 0.1 mm. (Fig. 2, Map 2).

#### RECOGNITION

Morphologically similar to *Inversodicraea liberia* Cheek in having 6-ribbed fruit capsules with weakly developed commissural ribs. However, *I. lebbiei* has narrowly triangular scale-leaves and not obovate scale-leaves as in *I. liberia*. Scale-leaves in *I. lebbiei* are acute at apex not broadly rounded as in *I. liberia* (Table 1. Moreover, *I. lebbiei* has 1 stamen, while *I. liberia* has 2 stamens.

#### SPECIMENS EXAMINED. SIERRA LEONE

Southern region, Kenema District, Sewa River, near Fomaya, Little Bekongor Falls, 8° 30’ 41.2” N; 11° 16’ 53.6” W, alt. 243 m, fl., fr., 3 May 2017, *Lebbie* 2709 (holotype K [K001971287]; isotypes MO, P, SL, Z). Southern region, Pujehun District, Waanje River, near Yawei Village, 7° 24’ 27.1” N; 11° 42’ 06.7” W, alt. 25 m, fl., fr., 11 May 2017, *Lebbie* 2726 (K [K001971288], SL, WAG).

#### CONSERVATION STATUS

The collection sites for *Inversodicraea lebbiei* are under anthropogenic threat. Artisanal mining for gold, agricultural runoff and domestic activities, such as clothes washing, threaten the long-term survival of this species. The collection locality in the Sewa River is at 243 m altitude and there may be suitable sites in this river for the construction of a hydroelectric dam, which poses a potential further/future threat.

*Inversodicraea lebbiei* is known from 2 collections made at 130 km from each other, in two distinct river systems which are 30 km distant or more from each other. According to IUCN (2024) two specimens cannot be used to calculate the extent of occurrence effectively. The area of occupancy is 8 km^2^. because the extent of occurrence must be equal to, or higher than, the area of occupancy (IUCN, 2024), the extent of occurrence is 8 km^2^. However, this species may also occur in some of the many other rapids in the two rivers. Given that some sites were not surveyed due to the curtailing of field work caused by rising water levels, a polygon of possible occurrence points suggests that the upper bound for the extent of occurrence may be between 1000 km^2^ and 4000 km^2^. The corresponding upper bound for area of occupancy may be between 8 km^2^ and 500 km^2^. The two collecting sites represent two locations, but because this species may also occur in other rapids, it is possible that the true number of locations is higher. The threat of dam construction is mainly located at Sewa River site, whiles the threats from gold mining, agricultural runoff and clothes washing occur in both locations. There exists a continuing decline in quality of habitat, and in number of mature individuals. *Inversodicraea lebbiei* is here preliminarily assessed as Endangered, EN B2ab(iii,v).

#### HABITAT

*Inversodicraea lebbiei* was collected on rocks in the Sewa River, 25 – 243 m alt., with *Tristicha trifaria* and *Letestuella* G. Taylor. The last was found mixed with *Inversodicraea lebbiei* in the collection *Lebbie, A*. 2709. The *Letestuella* sp. is now designated separately as *Lebbie* 2709A. This is the first record of *Letestuella* from Sierra Leone (POWO, continuously updated; gbif.org).

#### DISTRIBUTION

Sierra Leone, Southern Region, Kenema District and Pujehun District, Sewa River and Waanje River.

#### ETYMOLOGY

Named after Prof. Aiah Lebbie, Head of the National Herbarium of Sierra Leone (Njala University), now Vice-Chancellor & Principal of the University of Sierra Leone, who is the only collector of this species.

#### NOTES

*Inversodicraea lebbiei* has similarities with another remarkably distinct Sierra Leonian endemic, highly threatened species, *I. mortonii* (C.Cusset) Cheek (see Cheek *et al*. 2017a) in the triangular, stout, unlobed, scale-leaves. However, those of *I. lebbiei* are far shorter and broader than those of the last species which can be linear-conical. The presence in one collection of plants which seem annual, with short, slender stems fruiting directly from the root thallus, with other plants with similar stem borne on thick, almost woody naked branches c. 2 mm diam. which are possibly perennial, is remarkable. Species of the genus thus far are thought to be either annual (as in the first case) or perennial with longer, thicker stems, as in the second case. If later studies show that *I*.*lebbiei* is facultative in life-form, as seems likely, this will be a first in the genus.

## Discussion

The two newly identified species of *Inversodicraea, I. joulei*, and *I. lebbiei*, have contributed to the existing knowledge of the genus’s morphology and highlight the limited knowledge of Podostemaceae both in Sierra Leone and generally in Africa. Following the recent synoptic review of *Inversodicrea* (30 species) by Cheek *et al*. (2017a), eight additional species have been incorporated. Two species, *I. koukotamba* Cheek and *I. tassing* Cheek, from Guinea (Cheek *et al*. 2019), three Critically Endangered *I. senei* Cheek, *I. tanzaniensis* Cheek, and *I. botswana* Cheek from elsewhere in tropical Africa (Cheek *et al*. 2020); *I. nicolasii* (C.Cusset) E.Bidault, Rutish. & Mesterházy from Gabon (Bidault *et al*. 2023) and now these two from Sierra Leone result in a total of 38 species, making it the largest African genus in Podostemaceae for species diversity, now surpassing *Ledermanniella* which has 30 species,. Not only do these new species expand the known diversity of the genus, but they also support the evidence that Sierra Leone is a centre of endemism for Podostemaceae. The paper also presents the first taxonomic key specifically tailored to *Inversodicraea* species in Sierra Leone. Similar taxonomic work has been done in Cameroon (Cusset, 1987), but no such tool was previously available for Sierra Leone.

The study also reveals the presence of possible unidentified species among the rest of the collections from which the two new species were identified. Most of the Podostemaceae species in Sierra Leone are endemic or near endemic, including the near-endemic genus *Lebbiea*. However, it is hopeful that some of these ecosystems may be protected due to the Tropical Important Plant Areas (TIPAs) programme in Sierra Leone, an initiative that aims to discover and evidence regions with irreplaceable plant biodiversity so that they might be conserved against the risk of extinction (Darbyshire *et al*. 2017, Cheek *et al*. 2017a) following the model of neighbouring Guinea (Couch *et al*. 2019). Further systematic taxonomic work on Podostemaceae in Sierra Leone should be conducted after assessing further river sites fully since there are undoubtedly further species unknown to science that remain to be found.

## Acknowledgements

This paper was completed as part of a master’s project funded by B. A Krukoff bursary 2024 –25, Queen Mary University of London and Royal Kew Botanical Gardens ( Project ref. MScProject 2025 – 23). The authors thank the Joule Africa Project for facilitating the collection of these specimens for the Bumbuna Dam expansion. The National History Museum staff are appreciated for facilitating examination of specimens from their herbarium. Members of the Lebbie team, especially Momoh P. Sesay and Samuel O. Sokpo, are thanked for their unwavering dedication to collecting these specimens and their curation at the national herbarium of Sierra Leone.

## REFERENCES

Bachman, S., Moat, J., Hill, A. W., de la Torre, J. & Scott, B. (2011). Supporting Red List threat assessments with GeoCAT: Geospatial conservation assessment tool. ZooKeys, 150: 117–126. 10.3897/zookeys.150.2109

Bidault, E., Boupoya, A., Ikabanga, D. U., Nguimbit, I., Texier, N., Rutishauser, R., Mesterházy, A. & Stévart, T. (2023). Novitates Gabonenses 93. Plant Ecology and Evolution 156(1): 59–84.

Cheek, M. & Haba, P. M. (2016). Inversodicraea Engl. resurrected and I. pepehabai sp. nov. (Podostemaceae), a submontane forest species from the Republic of Guinea. Kew Bulletin 71: 55. 10.1007/S12225-016-9673-2.

Cheek, M. & Lebbie, A. (2018). Lebbiea (Podostemaceae-Podostemoideae), a new, nearly extinct genus with foliose tepals, in Sierra Leone. PLOS ONE, 13 (10): e0203603. 10.1371/journal.pone.0203603

Cheek, M., Feika, A., Lebbie, A., Goyder, D., Tchiengue, B., Sene, O., Tchouto, P. & van der Burgt, X. (2017a). A synoptic revision of Inversodicraea (Podostemaceae). Blumea -Biodiversity, Evolution and Biogeography of Plants, 62 (2): 125–156. 10.3767/blumea.2017.62.02.07

Cheek, M., Molmou, D., Jennings, L., Magassouba, S., & van der Burgt, X. (2019). Inversodicraea koukoutamba and I. tassing (Podostemaceae), new waterfall species from Guinea, West Africa. Blumea-Biodiversity, Evolution and Biogeography of Plants 64(3): 216–224.

Cheek, M., Molmou, D., Magassouba, S. & Ghogue, J.-P. (2022). Taxonomic revision of Saxicolella (Podostemaceae), African waterfall plants highly threatened by Hydro-Electric projects. Kew Bulletin 77 (2): 403–433. 10.1007/s12225-022-10019-2

Cheek, M., Séné, O., & Ngansop, E. (2020). Three new Critically Endangered Inversodicraea (Podostemaceae) species from Tropical Africa: I. senei, I. tanzaniensis and I. botswana. Kew Bulletin 75(2): 31.

Cheek, M., Van Der Burgt, X., Momoh, J. & Lebbie, A. (2017b). Ledermanniella yiben sp. nov. (Podostemaceae), Critically Endangered at the proposed Yiben Reservoir, Sierra Leone. Kew Bulletin 72 (2). 10.1007/s12225-017-9699-0

Couch, C., Cheek, M., Haba, P.M., Molmou, D., Williams, J., Magassouba, S., Doumbouya, S. & Diallo Y.M. (2019). Threatened habitats and Important Plant Areas (TIPAs) of Guinea, west Africa. Royal Botanic Gardens, Kew. London. https://kew.iro.bl.uk/concern/books/1554f509-3e22-453c-9fab-51b45722d250

Cusset, C. (1973). Contribution à l’étude des Podostemaceae. III. Le genre Stonesia. Bulletin Du Museum National d’Histoire Naturelle, Paris, 4e sér, 6, Sect. B, Adansonia, 13: 307–312. 10.5962/p.296772

Cusset, C. (1984). Contribution à l’étude des Podostemaceae. VIII: Ledermanniella Engl. sous-genre Ledermanniella. Bulletin Du Museum National d’Histoire Naturelle, Paris, 4e sér, 6, Sect. B, Adansonia, 3: 249–278.

Cusset, C. (1987). Podostemaceae. Flore Du Cameroon 30. MESRES, Yaoundé, Cameroon, 30: 51–101. https://www.nhbs.com/flore-du-cameroun-volume-30-book

Darbyshire, I., Anderson, S., Asatryan, A., Byfield, A., Cheek, M., Clubbe, C., Ghrabi, Z., Harris, T., Heatubun, C. D., Kalema, J., Magassouba, S., McCarthy, B., Milliken, W., de Montmollin, B., Lughadha, E. N., Onana, J.-M., Saïdou, D., Sârbu, A., Shrestha, K. & Radford, E. A. (2017). Important Plant Areas: Revised selection criteria for a global approach to plant conservation. Biodiversity and Conservation, 26 (8): 1767–1800. 10.1007/s10531-017-1336-6

Fayiah, M., Kallon, B. F., Dong, S., James, M. S. & Singh, S. (2020). Species Diversity, Growth, Status, and Biovolume of Taia River Riparian Forest in Southern Sierra Leone: Implications for Community-Based Conservation. International Journal of Forestry Research 2020 (1): 2198573. 10.1155/2020/2198573

IPNI. (continuously updated). International Plant Names Index. [Retrieved 27 May 2025]. https://Www.Ipni.Org/. https://www.ipni.org/search?q=Podostemaceae

IUCN (2012). IUCN Red List Categories and Criteria: Version 3.1. Second edition. IUCN, Gland, Cambridge.

IUCN (2024). IUCN Standards and Petitions Committee. Guidelines for Using the IUCN Red List Categories and Criteria. Version 16. Prepared by the Standards and Petitions Committee. https://www.iucnredlist.org/documents/RedListGuidelines.pdf. - Google Search

Kato, M. (2016). Multidisciplinary studies of diversity and evolution in riverweeds. Journal of Plant Research 129: 397–410. 10.1007/s10265-016-0801-8

Koi, S., Kita, Y., Hirayama, Y., Rutishauser, R., Huber, K. A. & Kato, M. (2012). Molecular phylogenetic analysis of Podostemaceae: Implications for taxonomy of major groups. Botanical Journal of the Linnean Society 169 (3): 461–492. 10.1111/j.1095-8339.2012.01258.x

Lebbie, A. (2017). A Survey of Ledermanniella yiben (Podostemaceae) in Selected River Rapids and Waterfalls in Sierra Leone. unpublished report, National Herbarium, Njala Univ., Sierra Leone. 19pp. https://selihydropower.sl/Content/documents/Ledermanniella%20yiben%20Field%20Report%202017%20Dr%20Lebbie%2020July17.pdf

Massally, F. K. (2025). A Taxonomic Overview of Podostemaceae (Orchids of the Falls) in Sierra Leone. Unpublished MSc thesis, Queen Mary University of London. https://kew.iro.bl.uk/concern/thesis_or_dissertations/db2c85ad-10d8-4fce-945c-1078118cc82a

Norman, P. E., Bebeley, J. F., Sesay, J. V. & Norman, Y. S. (2018). Biodiversity in Sierra Leone. In Global Biodiversity (1st Edition, pp. 213–246). Apple Academic Press

Philbrick, C. T. & Novelo, R. A. (1995). New World Podostemaceae: Ecological and evolutionary enigmas. Brittonia, 47 (2): 210–222. 10.2307/2806959

POWO. (continuously updated). Plants of the World Online. Retrieved 28 May 2025. https://Powo.Science.Kew.Org/. https://powo.science.kew.org/results?q=Podostemaceae

Rutishauser, R. (1995). Developmental patterns of leaves in Podostemaceae compared with more typical flowering plants: saltational evolution and fuzzy morphology. Canadian Journal Botany 73: 1305–1317.

Rutishauser, R., Novelo R A., & Philbrick, C. T. (1999). Developmental morphology of new world Podostemaceae: Marathrum and Vanroyenella. International Journal of Plant Sciences 160(1): 29–45.

Rutishauser, R. (2004). Podostemaceae of Africa and Madagascar: keys to genera and species, including genera description, illustrations to all species known, synonyms, and literature cited. https://www.systbot.uzh.ch/static/podostemaceae/keys/podostemaceae_key.pdf

Sierra Leone (2019). Residual Impact Assessment; Bumbuna II Hydropower Project. https://selihydropower.sl/Content/documents/Seli%20Residual%20Impact%20Assessment%202019.pdf

Thiv, M., Ghogue, J. P., Grob, V., Huber, K., Pfeifer, E., & Rutishauser, R. (2009). How to get off the mismatch at the generic rank in African Podostemaceae? Plant Systematics and Evolution 283: 57–77.

